# Loss of VHL-mediated pRb regulation promotes clear cell renal cell carcinoma

**DOI:** 10.1101/2024.04.14.589424

**Authors:** Mercy Akuma, Minjun Kim, Chenxuan Zhu, Ella Wiljer, Antoine Gaudreau-Lapierre, Leshan D. Patterson, Laura Trinkle-Mulcahy, William L. Stanford, Yasser Riazalhosseini, Ryan C. Russell

**Affiliations:** Department of Cellular and Molecular Medicine, University of Ottawa, Ottawa, ON, K1H 8M5, Canada; Department of Human Genetics, McGill University, Montreal, QC, H3A 0G1, Canada; Ottawa Hospital Research Institute (OHRI), Ottawa, ON, K1H 8L6, Canada; Department of Science, University of Waterloo, Waterloo, ON, N2L 3G1, Canada; Department of Biochemistry, University of Toronto, Toronto, Ontario, Canada; Ottawa Institute of Systems Biology, University of Ottawa, Canada; University of Ottawa Centre for Infection, Immunity and Inflammation, Ottawa, ON, Canada

**Author notes:** Corresponding author. Address for correspondence: 451 Smyth Rd, Ottawa, Ontario, K1H 8M5, Canada.

## Abstract

The von Hippel-Lindau (VHL) tumor suppressor is a component of E3 ubiquitin ligase complexes that target cellular substrates for proteasome-mediated degradation. VHL inactivation by genetic aberrations is observed in most sporadic cases of clear cell renal cell carcinoma (ccRCC). VHL loss leads to constitutive stabilization of E3 ligase targets, including hypoxia inducible factor α (HIFα), in VHL-associated tumors. HIFα stabilization upon VHL loss promotes transactivation of hypoxia responsive genes, which contributes to ccRCC development. However, several HIF-independent VHL targets have also been implicated in the promotion of tumorigenesis. Using proximity labeling to identify proteasomal VHL interactors, we identified retinoblastoma protein (pRb) as a novel substrate of VHL. Mechanistically, VHL interacts with pRb in an oxygen-sensitive manner, promoting its ubiquitin-mediated degradation. Concordantly, VHL-inactivation results in pRb hyperstabilization. Functionally, the hyperstabilization of pRb in ccRCC promoted tumorigenesis *in vitro* and in mouse models. We also show that downstream transcriptional changes induced by pRb hyperstabilization may contribute to ccRCC tumor development. Together, our findings reveal a novel VHL-related pathway which can be therapeutically targeted to inhibit ccRCC tumor development.

## Introduction

Von Hippel-Lindau (VHL) disease is a familial autosomal dominant disorder characterized by the predisposition to develop highly vascularized and glycolytic tumors in multiple organs^1^. VHL disease conforms to the Knudson’s ‘two-hit’ hypothesis^2–4^. Germline mutations in VHL result in a defective allele and this represents the first ‘hit’. The second ‘hit’ is a somatic event which occurs in the second allele, leading to loss of heterozygosity at the VHL locus. Inactivation of both VHL alleles can lead to tumor development in the affected tissue. The clinical manifestations of VHL disease include retinal and central nervous system hemangioblastomas, clear cell renal cell carcinoma (ccRCC), pheochromocytomas, pancreatic neuroendocrine tumors, and endolymphatic sac tumors (ELSTs)^1,5–10^. However, the leading cause of mortality in patients with VHL disease is metastatic ccRCC^11^.

Biallelic VHL inactivation has also been observed in about 70% of sporadic ccRCCs^12,13^. The best characterized function of VHL relates to its role as the substrate recognition component of an E3 ubiquitin ligase complex that also contains Elongin B, Elongin C, Cullin-2 and Rbx1 (called VCB-CUL2 complex)^14–16^. The VCB-CUL2 complex promotes the ubiquitination and proteasomal degradation of the hypoxia inducible factor α (HIFα) under normoxia^17,18^. VHL inactivation in ccRCC leads to constitutive HIFα stabilization, irrespective of oxygen levels^19^. HIFα activates genes that promote tumor adaptation and development, linking HIF stabilization to tumorigenesis^20–25^. Downregulation of HIF2α was shown to be sufficient to suppress tumorigenesis in VHL-defective ccRCC cells, whereas stable expression of a HIF2α variant lacking VHL recognition can override VHL-mediated tumor suppression in ccRCC cells^26,27^. As such, significant effort has been put into pharmacologically targeting HIF or HIF-related pathways^28–34^. However, the most significant therapeutic progress has been a result of combination treatments involving immune-oncologic drugs^35–37^. Despite improved response with the addition of immune-oncologic drugs, the majority of patients with metastatic ccRCC are still resistant to all current therapeutic options^38,39^. Therefore, identifying novel pathways underlying ccRCC is needed to establish new therapeutic targets.

There are several ‘non-HIF’ proteolytic substrates of VHL such as Epidermal Growth Factor Receptor (EGFR), Zinc finger and homeobox 2 (ZHX2), and β_2_-adrenergic receptor (β_2_AR) that are involved in signal transduction^40–42^. Other targets include atypical protein kinase C (aPKC), Sprouty2 (Spry2), and Scm-like with four malignant brain tumor domains 1 (SFMBT1), which regulate cellular differentiation^43–45^. These HIF-independent VHL substrates may also be involved in cancer development. For example, ZHX2 may promote ccRCC oncogenesis via activation of the Nuclear factor kappa B (NF-κB) pathway^41^. Also, SFMBT1, a key regulator of epithelial-to-mesenchymal transition, has been shown to be upregulated in ccRCC and contribute to tumor growth^45,46^. In support for the biological relevance of HIF-independent VHL targets, a synthetic mutant that retains HIF degradation, but is deficient in the regulation of some HIF-independent targets, was found to be incapable of suppressing ccRCC tumorigenesis^47,48^. These findings highlight the potential contributions of non-HIF substrates of VHL to tumor development. Interestingly, conditional knockout of pRb was shown to be synthetically lethal with VHL loss in the retina, another tissue impacted in VHL kindred^49^. The increase in cell death in *Vhl/Rb1* double knockout retinas was also accompanied by increased angiogenesis in unaffected cells indicating an abnormal response to the pseudohypoxia caused by *Vhl* loss. Additionally, *Rb1* deletion in the mouse fetal liver resulted in an increase in cell death in hypoxic tissue^50^. Together, these studies indicate that VHL and pRb may both be involved in regulating cellular viability in response to hypoxia.

pRb is involved in the regulation of cellular processes such as G1-S transition, DNA damage checkpoint, cell cycle exit, and cellular differentiation^51–54^. When hypophosphorylated, pRb interacts with the E2F transcription factors, inhibiting the transcription of genes required for S-phase entry^55–59^. However, when the cell is ready to divide, pRb becomes hyperphosphorylated, inhibiting its ability to sequester E2Fs and thereby promoting cell cycle. Hence, the original tumor suppressive function described for pRb relates to its ability to inhibit cell cycle progression in its hypophosphorylated state. It is therefore common for tumor cells to inactivate pRb’s growth suppressive function by exploiting pathways that regulate pRb phosphorylation^60,61^. For example, cyclin D1 overexpression observed in ccRCC cells results in pRb hyperphosphorylation, which supports abnormal proliferation in cells deprived of growth factors^62^.

pRb has also been shown to regulate cell-cycle-independent pathways such as inflammation, autophagy, apoptosis, metabolism and stemness^63–68^, which have the potential to contribute to oncogenesis. For instance, pRb has been described to have anti-apoptotic functions and its inactivation is associated with increased levels of apoptosis^69–72^. pRb has also been shown to inhibit hypoxia-mediated induction of angiogenesis and autophagic cell death, as mentioned previously^49,50^. Hence, pRb may have the capacity to support oncogenesis depending on the context. Accordingly, pRb overexpression has been observed in multiple cancer types including familial adrenocortical carcinomas and pancreatic adenocarcinomas^73,74^. Furthermore, transgene expression of phosphorylation-resistant constitutively active pRb in the mammary gland led to the development of mammary adenocarcinoma^75^.

In this study, we characterize pRb as a novel substrate of VHL, which is targeted for ubiquitin-dependent degradation in an oxygen-sensitive manner. We found that pRb was significantly upregulated in ccRCC and that high pRb protein expression is associated with increased tumorigenic potential of ccRCC cells. We further demonstrate that pRb inhibits apoptosis in ccRCC cells and that this function may be via regulation of downstream transcriptional targets including Ski/Dach domain-containing protein 1 (SKIDA1), a minimally characterized protein. Overall, our findings highlight the oxygen-sensitive regulation of pRb by VHL and the potential implications of VHL-pRb dysregulation on hypoxia-mediated cell death in ccRCC tumors. Importantly, we present a novel VHL-related pathway which can be therapeutically targeted to inhibit ccRCC tumor development.

## Results

### pRb binds VHL in a proteasomal-sensitive manner

The interaction between VHL and HIF is stabilized in the presence of proteasome inhibitors^76–78^. To identify novel proteasome-sensitive VHL substrates that might contribute to disease, we performed a proximity-dependent biotin identification (BioID) assay using HEK293A cells expressing VHL fused to the biotin ligase BirA. Cells were pre-treated with the proteasome inhibitor MG132 or vehicle control for 4 hours followed by 1 hour of biotin addition. MG132 stabilizes ubiquitinated proteins marked for degradation by blocking the proteolytic activity of the 26S proteasome complex and this is known to stabilize the interaction of VHL and its targets, including HIFα^19^. Biotinylated proteins were captured using streptavidin-conjugated beads and resolved on an SDS-PAGE gel. Proteins were visualized following a Coomassie stain. Protein-containing gel slices were then excised, trypsin digested, and analyzed by liquid chromatography-mass spectrometry (LC-MS) (Figure 1A). LC–MS analysis revealed several proteins that were enriched in MG132-treated cells compared to vehicle control. Interestingly, the tumor suppressor pRb was one of the proteins enriched following MG132 treatment, raising the possibility that pRb is a proteasomal target of VHL. Despite the apparent paradox of one tumor suppressor degrading another, the prior data linking *Vhl* and *Rb1* in synthetic lethality and pRb’s role in modulating hypoxic cell death led us to validate pRb as our top hit.

**Figure 1.**
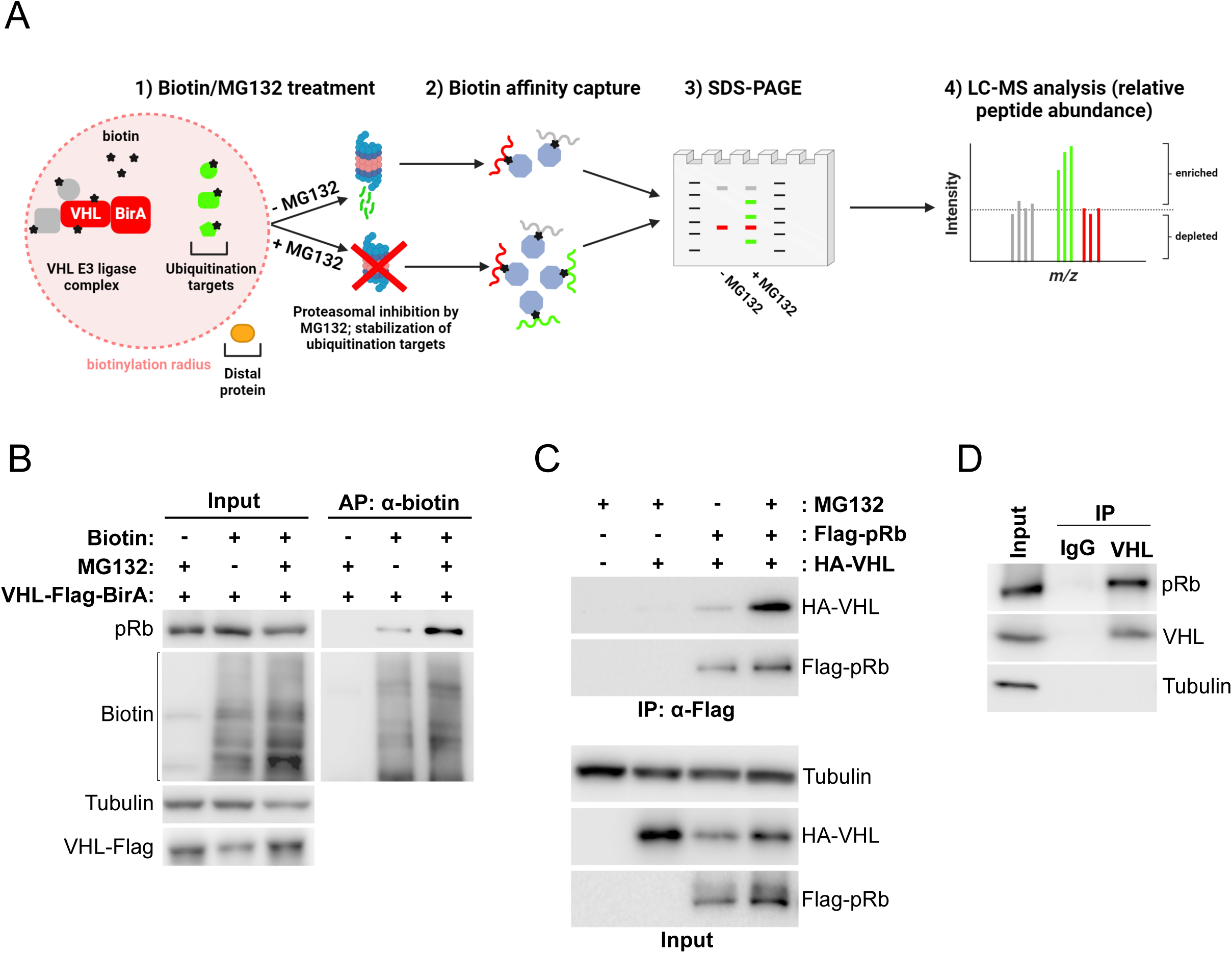
Proximity ligation analysis of VHL reveals pRb as a proteasome-sensitive VHL interactor. A. Schematic diagram illustrating BioID assay used to identify VHL interacting proteins. Figure was created with BioRender.com. In the experimental condition, 10 µM MG132 was added for 4 h, followed by 50 µM biotin for 1 h. In the control condition, 10 µM DMSO vehicle was added instead of MG132. Biotinylated proteins were captured using streptavidin-conjugated beads and resolved on SDS-PAGE gel. Gel chunks were then extracted separately from control and experimental lanes, trypsin digested, and analysed by mass spectrometry. B. Immunoprecipitation of biotinylated proteins using streptavidin-conjugated beads from HEK293A cells transfected with VHL-Flag-BirA plasmid and treated with 10 µM MG132 and 50 µM biotin as indicated for 4 h and 1 h respectively. C. Immunoprecipitation of Flag-tagged pRb from HEK293A cells transfected with indicated plasmids and treated with 10 µM MG132 as indicated for 4 h. D. Immunoprecipitation of endogenous VHL from HEK293A cells treated with 10 µM MG132 for 4 h. An IgG1 isotype control was included. MG132-treated HEK293A lysate was split equally between IgG1 and VHL pulldown conditions.

To validate our mass spectrometry results, we first sought to confirm the enrichment of pRb in samples preferentially biotinylated by VHL-BirA in MG132-treated cells. Cells were transfected with VHL-BirA and treated with biotin and MG132 or vehicle as described above. Biotinylated proteins were pulled down with streptavidin beads and eluted proteins were resolved by SDS-PAGE. Immunoblot for pRb in the eluent showed a significant stabilization of VHL interaction with pRb following proteasomal blockade, consistent with our mass spectrometry data (Figure 1B). To determine if VHL and pRb interact, we exogenously expressed Flag-tagged pRb and HA-tagged VHL in HEK293A cells in the presence or absence of MG132. Immunoprecipitation of pRb revealed increased co-precipitation of VHL in the presence of MG132 verifying a proteasomal-sensitive interaction (Figure 1C). To determine if VHL and pRb interact endogenously, we performed immunoprecipitation on MG132-treated HEK293A lysates using either an anti-VHL or a non-targeting IgG antibody. Immunoblot analysis showed co-precipitation of pRb with endogenous VHL (Figure 1D). Together, these findings indicate that pRb is a novel binding partner of VHL that may be targeted by the ubiquitin-proteasome pathway, similar to HIFα.

### VHL regulates pRb stability via the ubiquitin-proteasome pathway

To determine the effect of VHL on pRb expression, HA-tagged VHL was reconstituted into two VHL-deficient ccRCC cell lines, 786-O and RCC4. Cells were lysed at 24 hours post-confluence to mitigate cell cycle variation in log-phase growth and any associated variability in pRb regulation by cell cycle dependent factors. Western blot analysis in both ccRCC lines showed that VHL re-expression leads to downregulation of pRb protein levels (Figure 2A). This effect was determined to be post-transcriptional, as pRb mRNA levels were not affected by VHL reconstitution, as measured by quantitative real-time PCR (qRT-PCR) (Figure 2B). Next, we sought to determine the impact of endogenous VHL knockdown in HEK293A and U2OS cells, which have functional VHL expression. In both cell lines, knockdown of VHL led to an increase in pRb protein expression (Figure 2C). Consistent with our results in the ccRCC cell lines, VHL status did not affect pRb mRNA levels (Figure 2D). Collectively, these results indicate that VHL regulates pRb at the post-transcriptional level.

**Figure 2.**
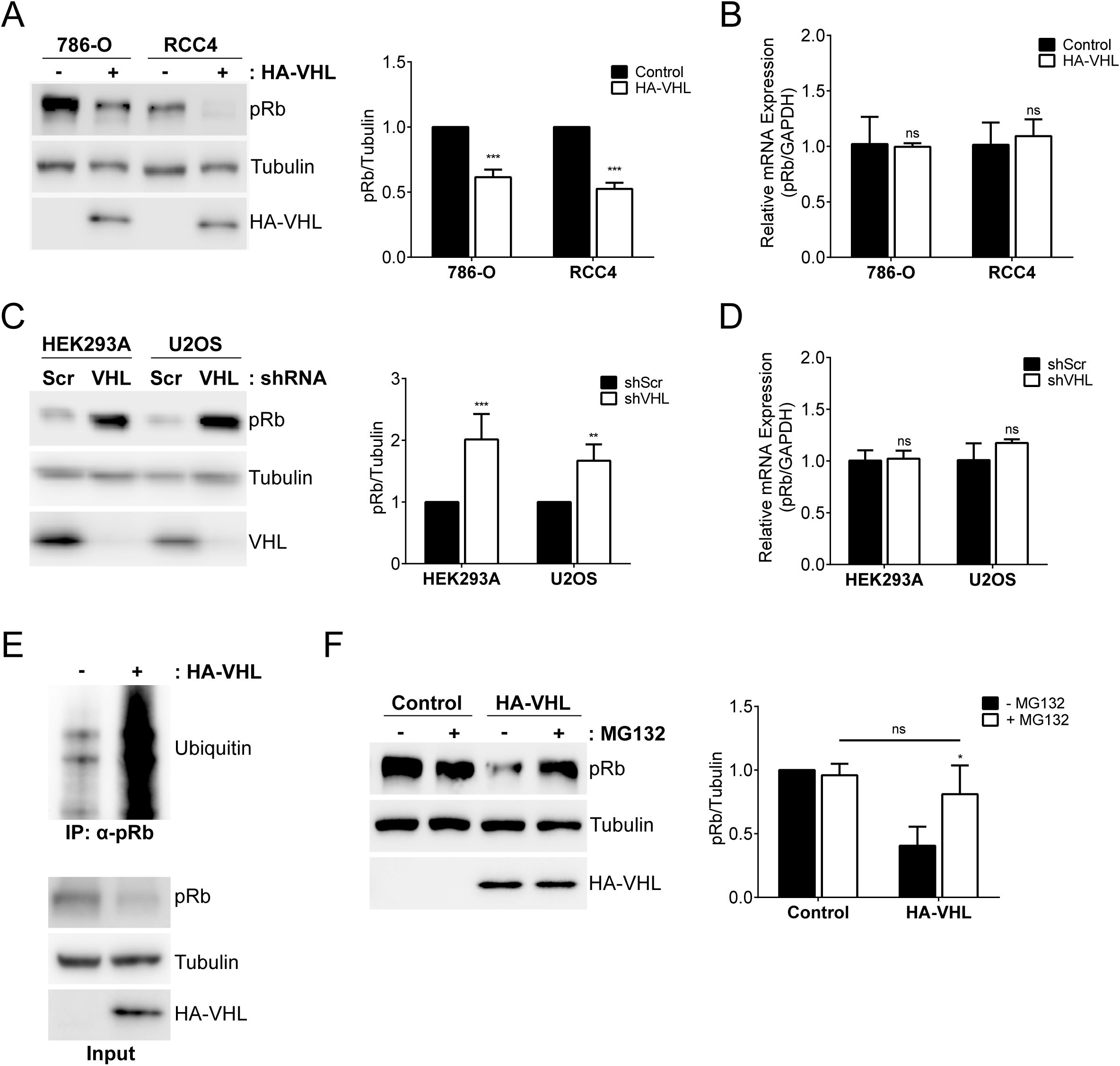
VHL promotes proteasomal degradation of pRb. A. Immunoblot of lysates from 786-O and RCC4 cells stably transfected with either control vector (-) or hemagglutinin (HA)-tagged VHL. Quantification of pRb protein expression relative to tubulin is shown to the right. Statistical significance was calculated using two-way ANOVA and Fisher’s LSD test. B. pRb/GAPDH mRNA expression in 786-O and RCC4 cells stably transfected with either control vector or HA-tagged VHL. Fold changes in gene expression were calculated using the delta delta Ct method. Statistical significance was calculated using two-way ANOVA and Fisher’s LSD test. C. Immunoblot of lysates from HEK293A and U2OS infected with lentivirus encoding either scrambled short hairpin RNA (shRNA) control vector (Scr) or VHL targeting shRNA. Quantification of pRb protein expression relative to tubulin is shown to the right. Statistical significance was calculated using two-way ANOVA and Fisher’s LSD test. D. pRb/GAPDH mRNA expression in HEK293A and U2OS cells infected with lentivirus encoding either scrambled (Scr) or VHL shRNA. Fold changes in gene expression were calculated using the delta delta Ct method. Statistical significance was calculated using two-way ANOVA and Fisher’s LSD test. E. Immunoprecipitation of denatured pRb from 786-O cells stably transfected with either control vector (-) or hemagglutinin (HA)-tagged VHL and treated with 10 µM MG132 for 4 h. F. Immunoblot of lysates from 786-O cells stably transfected with either control vector or HA-tagged VHL and treated as indicated with 10 µM MG132 for 4 h. Quantification of pRb protein expression relative to tubulin is shown to the right. Statistical significance was calculated using two-way ANOVA and Tukey’s HSD test. (A-F) * p < 0.05, ** p < 0.01, *** p < 0.001. ‘ns’ denotes not significant. Error bars represent standard deviation.

Given that VHL interaction with pRb is stabilized by proteasomal inhibition, we next asked whether VHL could promote ubiquitination of pRb. To analyze VHL-mediated ubiquitination of pRb, we immunoprecipitated pRb under denaturing conditions from VHL-null and VHL-reconstituted 786-O cells treated with MG132. Immunoblot analysis showed significant pRb ubiquitination only in cells expressing VHL (Figure 2E). Treatment of VHL-reconstituted 786-O cells with MG132 resulted in pRb protein accumulation comparable to VHL-null 786-O cells, whereas MG132 treatment of VHL-null cells did not significantly affect pRb protein levels (Figure 2F) Together, these data demonstrate that VHL promotes the ubiquitination of pRb, thereby facilitating its proteasomal degradation.

### Transcriptional regulation by the VHL-pRb and VHL-HIF pathways are largely distinct

The oxygen-dependent regulation of HIFα stability is important in the transcriptional reprogramming of cells under hypoxia^79^. pRb has also been described to repress hypoxia-regulated transcription^50^. However, the mechanism for oxygen-dependent pRb transcriptional regulation is not clearly understood. This led us to investigate whether VHL regulation of pRb was oxygen-sensitive. We analyzed the oxygen sensitivity of the pRb-VHL interaction in transfected cells. Immunoprecipitation of VHL showed that binding to pRb is inhibited by hypoxia (0.5% O_2_), demonstrating that VHL regulation of pRb is oxygen-dependent (Figure 3A). This suggests that pRb is involved in the cellular adaptation to hypoxia.

**Figure 3.**
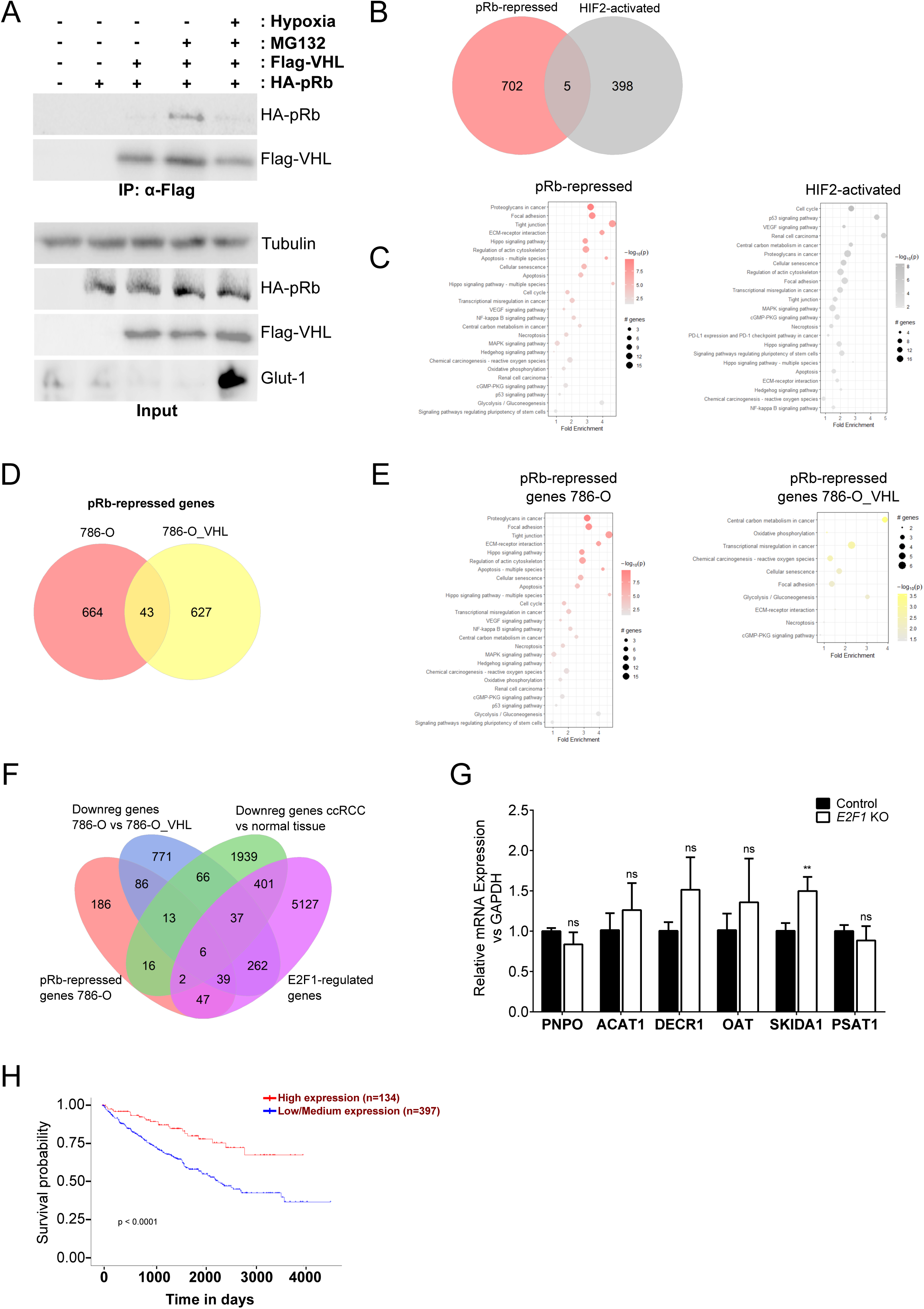
Transcriptional regulation by the VHL-pRb and VHL-HIF axes are distinct. A. Immunoprecipitation of Flag-tagged VHL from HEK293A cells transfected with indicated plasmids and treated with 10 µM MG132 as indicated for 4 h. For hypoxia treatment, cells were placed in 0.5% O_2_ for 24 h. Glut-1 upregulation serves as positive control to indicate oxygen depletion under hypoxia. B. Venn diagram showing overlap between genes upregulated by *RB1* KO in 786-O cells (red), and genes activated by HIF2α (grey). FC > 1.5, adjusted p-value < 0.05. C. KEGG pathway enrichment analysis of pRb-repressed and HIF-activated genes in 786-O cells D. Venn diagram showing overlap between genes upregulated by *RB1* KO in VHL-null (red) and VHL-reconstituted (yellow) 786-O cells. FC > 1.5, adjusted p-value < 0.05. E. KEGG pathway enrichment analysis of pRb-repressed genes in VHL-null and VHL-reconstituted 786-O cells F. Four-way Venn diagram showing overlap between the indicated gene sets. FC > 2, adjusted p-value < 0.05. E2F1-regulated transcripts were obtained from TRANSFAC database from ChIP-X enrichment analysis^82,108^. Downregulated transcripts in ccRCC vs normal tissue were obtained from the International Cancer Genome Consortium (ICGC) data portal^12^. Prognostic markers of ccRCC were obtained from the University of ALabama at Birmingham CANcer (UALCAN)^85^ data analysis Portal and based on Kaplan-Meier survival analysis^83^. G. Analysis of mRNA expression of indicated genes in 786-O Cas9-expressing cells stably infected with virus encoding either control vector or *E2F1*-targeting guides. Statistical significance was calculated using unpaired t-test. H. Kaplan-Meier analysis showing the correlation between SKIDA1 expression in ccRCC tumors and patient survival. (A-H) * p < 0.05, ** p < 0.01, *** p < 0.001. ‘ns’ denotes not significant. Error bars represent standard deviation.

In response to oxygen limitation, cells upregulate pathways such as angiogenesis, glucose metabolism and erythropoiesis to promote cellular adaptation to hypoxia^80^. pRb has also been described to regulate HIF-responsive genes in an oxygen-sensitive manner^49,50^. We first asked whether pRb and HIF coordinate to transcriptionally regulate the same target genes. To identify pRb-regulated genes in ccRCC, we first created a monoclonal *RB1* knockout (KO) cell line from 786-O cells using CRISPR/Cas9. We next determined global mRNA expression in 786-O control and *RB1* KO cells using RNA-sequencing. We were interested in genes that were upregulated following *RB1* KO, as pRb is known to act as a transcriptional repressor. We found 707 genes with significantly higher expression in *RB1* KO cells compared to control cells (fold change > 1.5, adjusted p-value (q) < 0.05) (Figure 3B, Supplementary Table S1). In contrast to pRb, HIF is a transcriptional activator. HIF2α target genes were obtained from published datasets of RNA-sequencing analysis of 786-O control and *EPAS1* (HIF2α) KO in 786-O cells^81^ (fold change > 1.5, q < 0.05). Analysis of pRb-repressed and HIF2α-activated genes showed little overlap, indicating that pRb regulates a distinct set of genes from HIF2α (Figure 3B). Despite having few common targets at the level of individual genes, KEGG pathway analysis of HIF and pRb-regulated gene sets showed enrichment of similar pathways such as focal adhesion, extracellular matrix (ECM)-receptor interaction, cell cycle, cellular senescence, and apoptosis (Figure 3C). These results indicate that the transcriptional influence of VHL deletion in ccRCC is impacted by a combined dysfunction in HIF2α and pRb transcriptional regulation.

To determine pRb-regulated transcription in low vs high HIF2α-expressing cells, we created *RB1* knockouts in both VHL-reconstituted and VHL-null 786-O cells (Supplementary Figure S1).

Following RNA-sequencing, we analyzed transcripts that were significantly upregulated in *RB1* KO cells compared to corresponding controls, to identify pRb-repressed genes. We identified 670 genes that were significantly repressed by pRb in VHL-reconstituted cells (Supplementary Table S2), compared to the 707 genes repressed in VHL-null 786-O cells as described above (fold change > 1.5, q < 0.05). Interestingly, comparison of both gene sets showed roughly 6% overlap, suggesting that pRb transcriptionally regulates a distinct set of genes based on HIF expression (Figure 3D). However, KEGG pathway analysis of pRb-repressed transcripts in VHL-null and VHL-reconstituted cell lines showed enrichment of similar pathways such as focal adhesion, ECM-receptor interaction, necroptosis, oxidative phosphorylation, glycolysis, and cellular senescence (Figure 3E). These tumor-associated pathways raise the possibility of the involvement of pRb in ccRCC oncogenesis.

We next sought to identify which transcriptional targets of pRb contribute to ccRCC development. To narrow down the list of potential candidates, a stricter selection criteria of fold change > 2 and q < 0.05 was applied. We omitted the pRb-repressed genes in cells with low HIF and VHL reconstitution, which differs from the conditions in most ccRCC cases, leaving 395 genes (Figure 3F, Supplementary Tables S1 and S2). We then removed genes whose expression were not sensitive to VHL status (251 genes removed, Supplementary Table S3), leaving 144 genes. Among these VHL and pRb differentially expressed genes, we removed any that were not regulated by E2F-1 (obtained from ChIP-X Enrichment Analysis datasets^82^), as pRb has been described to repress apoptosis largely through E2F1^54^, leaving 45 genes. Finally, we asked which of these genes were repressed in ccRCC tumors versus normal tissue (Supplementary Table S4), which resulted in 6 genes (PNPO, ACAT1, DECR1, OAT, SKIDA1 and PSAT1) that fulfilled the following criteria: (1) repressed by pRb in 786-O cells (2) downregulated in VHL-deficient vs VHL-reconstituted 786-O cells (3) downregulated in ccRCC tumor vs normal tissue, and (4) regulated by E2F1 (Figure 3F).

Among the selection criteria described above to identify physiological transcriptional targets of the VHL-pRb pathway, we have not demonstrated a requirement for E2F1 in our cell lines. To determine if the 6 narrowed down genes were regulated by E2F1 in ccRCC, we knocked out E2F1 from 786-O cells using CRISPR/Cas9 (Supplementary Figure S2) and examined the effect on mRNA expression of each gene. qRT-PCR analysis demonstrated that SKI/DACH domain containing protein 1 (SKIDA1) is transcriptionally repressed in ccRCC cells in an E2F1-dependent manner (Figure 3G). DECR1 and OAT mRNA levels were also increased upon E2F1 KO, but did not reach statistical significance. We next confirmed that SKIDA1 expression was upregulated by *RB1* deletion by qRT-PCR and immunoblot analysis of lysates from 786-O control and *RB1* KO cells (Supplementary Figures S3 and S4). These results suggest that SKIDA1 may be a downstream target of the VHL-pRb pathway and may be functionally repressed in ccRCC patients. Kaplan-Meier^83^ survival analysis of ccRCC patients revealed a positive correlation between SKIDA1 expression in tumors and patient survival (Figure 3H). These findings highlight SKIDA1 as a putative pRb target with prognostic significance in ccRCC, and whose repression upon VHL deletion may contribute to tumorigenesis.

### pRb inhibits apoptosis in ccRCC cells

We have demonstrated that VHL promotes the degradation of pRb in an oxygen and proteasomal sensitive manner. Therefore, pRb may be hyperstabilized in VHL-deficient tumors. Proteomic data obtained from the Clinical Proteomic Tumor Analysis Consortium (CPTAC)^84,85^ confirmed that pRb protein abundance is higher in ccRCC primary tumors (n=110) compared to normal tissue (n=84) (Figure 4A). Consistent with CPTAC data, immunoblot analysis showed that pRb was upregulated in the majority of ccRCC tumors compared to patient-matched normal tissue (Figure 4B). These patient-derived data are consistent with our *in vitro* work in ccRCC cells, which showed a dramatic increase of pRb levels in cells lacking functional VHL (Figure 2A).

**Figure 4.**
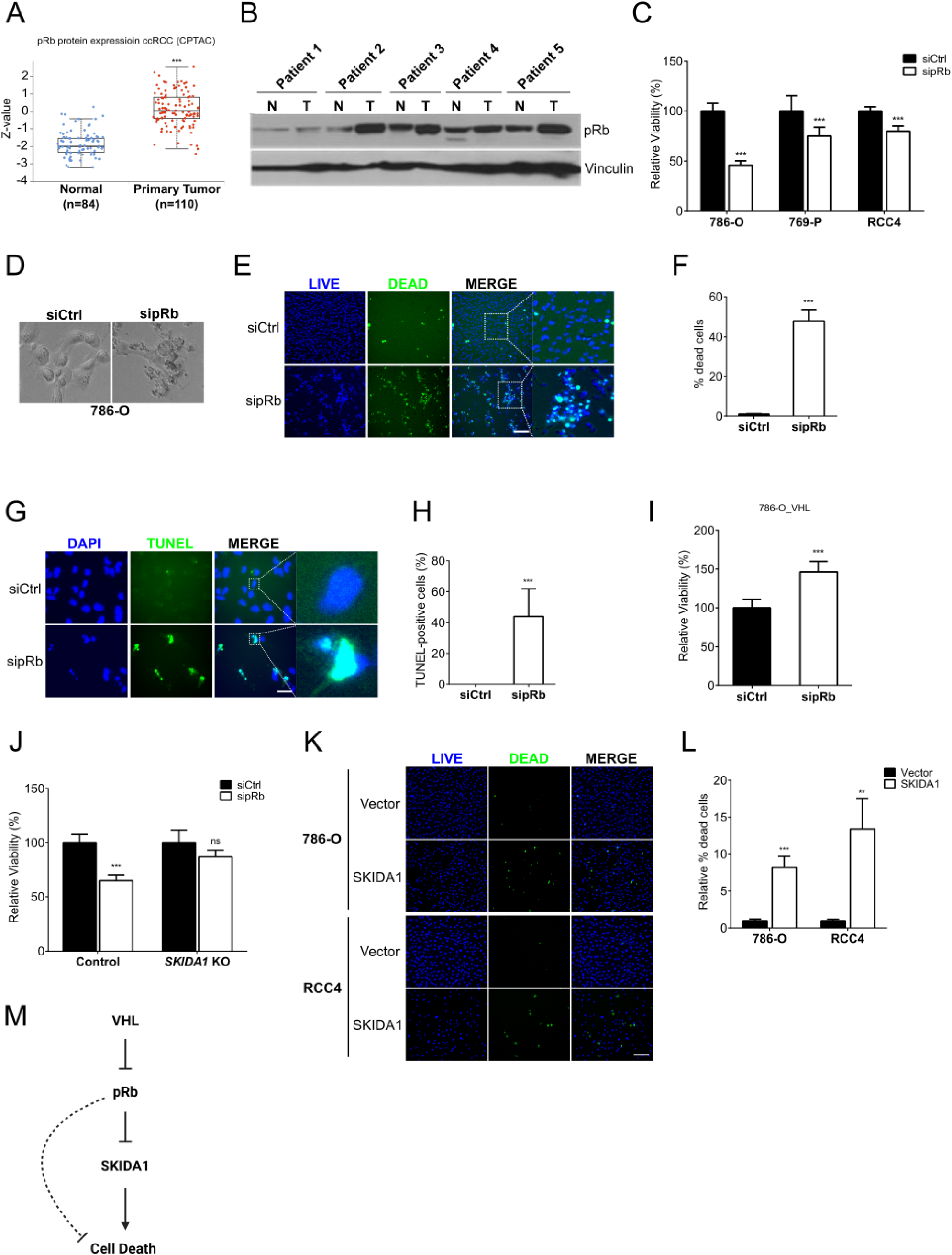
pRb suppresses apoptosis in ccRCC cells. A. Analysis of pRb protein expression in ccRCC primary tumors compared to normal tissue. Patient data were obtained from Clinical Proteomic Tumor Analysis Consortium (CPTAC) dataset via the UALCAN data analysis portal. Z-values represent standard deviations from the median across normal and tumor samples. Log2 spectral count ratio values from CPTAC were first normalized within each sample profile, then normalized across samples. B. Immunoblot analysis of lysates from ccRCC primary tumor (T) or patient-matched normal tissue (N). C. Analysis of cell viability in indicated ccRCC cell lines transfected with either non-targeting control siRNA (siCtrl) or pRb targeting siRNA (sipRb), measured via a sulforhodamine B (SRB) assay. Statistical significance was calculated using unpaired t-test. D. Representative phase-contrast images of 786-O cells transfected with either non-targeting control siRNA (siCtrl) or pRb targeting siRNA (sipRb) E. Representative fluorescence images of 786-O cells transfected with either non-targeting control siRNA (siCtrl) or pRb targeting siRNA (sipRb), and stained using the ReadyProbes® Cell Viability Imaging Kit. The blue stain represents total cells, while the green stain represents dead cells. Scale bar = 250 µm. F. Quantification of percentage of dead cells relative to total number of cells in (F). Statistical significance was calculated using unpaired t-test. G. Representative TUNEL staining of 786-O cells transfected with either non-targeting control siRNA (siCtrl) or pRb targeting siRNA (sipRb). Scale bar = 250 µm. H. Quantification of percentage of TUNEL-positive cells relative to total number of cells in (H). Statistical significance was calculated using unpaired t-test. I. Analysis of cell viability in VHL-expressing 786-O cells transfected with either non-targeting control siRNA (siCtrl) or pRb targeting siRNA (sipRb), measured via a sulforhodamine B (SRB) assay. Statistical significance was calculated using unpaired t-test. J. Analysis of cell viability in 786-O control or *SKIDA1* KO cells, transfected with either non-targeting control siRNA (siCtrl) or pRb targeting siRNA (sipRb), measured via a sulforhodamine B (SRB) assay. Statistical significance was calculated using two-way ANOVA and Fisher’s LSD test. K. Representative fluorescence images of 786-O and RCC4 cells transduced with lentivirus encoding either control plasmid (Vector) or SKIDA1 cDNA containing plasmid. Cells were stained using the ReadyProbes® Cell Viability Imaging Kit. The blue stain represents total cells, while the green stain represents dead cells. Scale bar = 250 µm. L. Quantification of percentage of dead cells relative to total number of cells in (K). Statistical significance was calculated using unpaired t-test. M. Schematic illustration showing the VHL-pRb-SKIDA1 signalling axis and effect on cell death. The dotted line indicates indirect regulation. (A-M) * p < 0.05, ** p < 0.01, *** p < 0.001. ‘ns’ denotes not significant. Error bars represent standard deviation.

To determine the effect of pRb hyperstabilization on viability in ccRCC cells, we depleted pRb from several VHL-deficient ccRCC cell lines using pRb-targeting siRNAs (Supplementary Figure S5) and analyzed cell viability using a sulforhodamine B (SRB) colorimetric assay. We observed a significant decrease in cell viability following pRb depletion in all ccRCC cell lines (Figure 4C). By microscopy, we observed morphological changes associated with cell death such as cell shrinkage, rounding, and increased cellular debris in pRb-depleted 786-O cells (Figure 4D). Live/dead cell staining of 786-O cells confirmed an increase in the proportion of dead cells upon siRNA-mediated knockdown of pRb (Figure 4E-F). Several reports have shown that pRb can repress apoptosis, while others have reported a repression of non-apoptotic cell death^50,54,72,86–90^. To determine if cell death induced by pRb depletion was apoptotic, we performed terminal deoxynucleotidyl transferase dUTP nick end labeling (TUNEL) staining on 786-O control and pRb knockdown cells. We observed a significant increase in the proportion of TUNEL-positive (or apoptotic) cells following pRb knockdown, suggesting that pRb inhibits apoptosis in ccRCC cells (Figure 4G-H). Interestingly, pRb knockdown in VHL-reconstituted 786-O cells did not lead to a similar decrease in cell viability, in stark contrast to the effect of pRb depletion in VHL-null cells (Figure 4I, Supplementary Figure S6). These findings suggest that the anti-apoptotic function of pRb in ccRCC cells may be dependent on VHL or HIF expression, which is consistent with the differential transcriptional regulation by pRb in the presence or absence of VHL (Figure 3D) and the synthetic lethality observed between VHL and *RB1*^49^.

We next asked whether the downstream VHL-pRb transcriptional target SKIDA1 was functionally significant in the induction of cell death following pRb knockdown. Since SKIDA1 is upregulated upon knockout of *RB1*, we knocked out *SKIDA1* in 786-O cells and analyzed the effect of pRb depletion on cell viability as measured by an SRB assay. Knockout of *SKIDA1* rescued viability of 786-O cells depleted of pRb (Figure 4J). Next, we tested if overexpression of SKIDA1 alone was sufficient to induce cell death. Two ccRCC cell lines were infected with lentivirus encoding either control vector or SKIDA1 expression vector (Supplementary Figure S7). Live/dead cell staining showed a significant increase in the proportion of dead cells following SKIDA1 overexpression in both cell lines (Figure 4K-L). Collectively, our results suggest that pRb suppresses cell death specifically in VHL-deficient renal cells, likely via regulation of downstream transcriptional targets such as SKIDA1 (Figure 4M).

### pRb depletion inhibits ccRCC tumorigenesis

pRb-mediated repression of cell death in ccRCC may be indicative of oncogenic contribution of hyperaccumulated pRb in ccRCC. Therefore, we sought to determine the effects of pRb accumulation on ccRCC cancer phenotype. We first analyzed anchorage-independent growth of ccRCC cells in soft agar, which is an important hallmark of cancer^91–93^. 786-O cells are known to grow in an anchorage-independent manner^41,81^, so in this background we created two independent *RB1* KO cell lines using CRISPR/Cas9 (Supplementary Figures S8 and S9). Next, using 786-O control and *RB1* KO cells, we measured anchorage-independent growth in the soft agar 3D culture model. We found that knockout of *RB1* significantly inhibited anchorage-independent growth of 786-O cells (Figure 5A-B). In contrast, knockout of *RB1* in VHL-reconstituted 786-O cells was sufficient to induce anchorage-independent growth (Figure 5C-D). These data are consistent with an oncogenic role for hyperaccumulated pRb specifically in VHL-deficient ccRCC.

**Figure 5.**
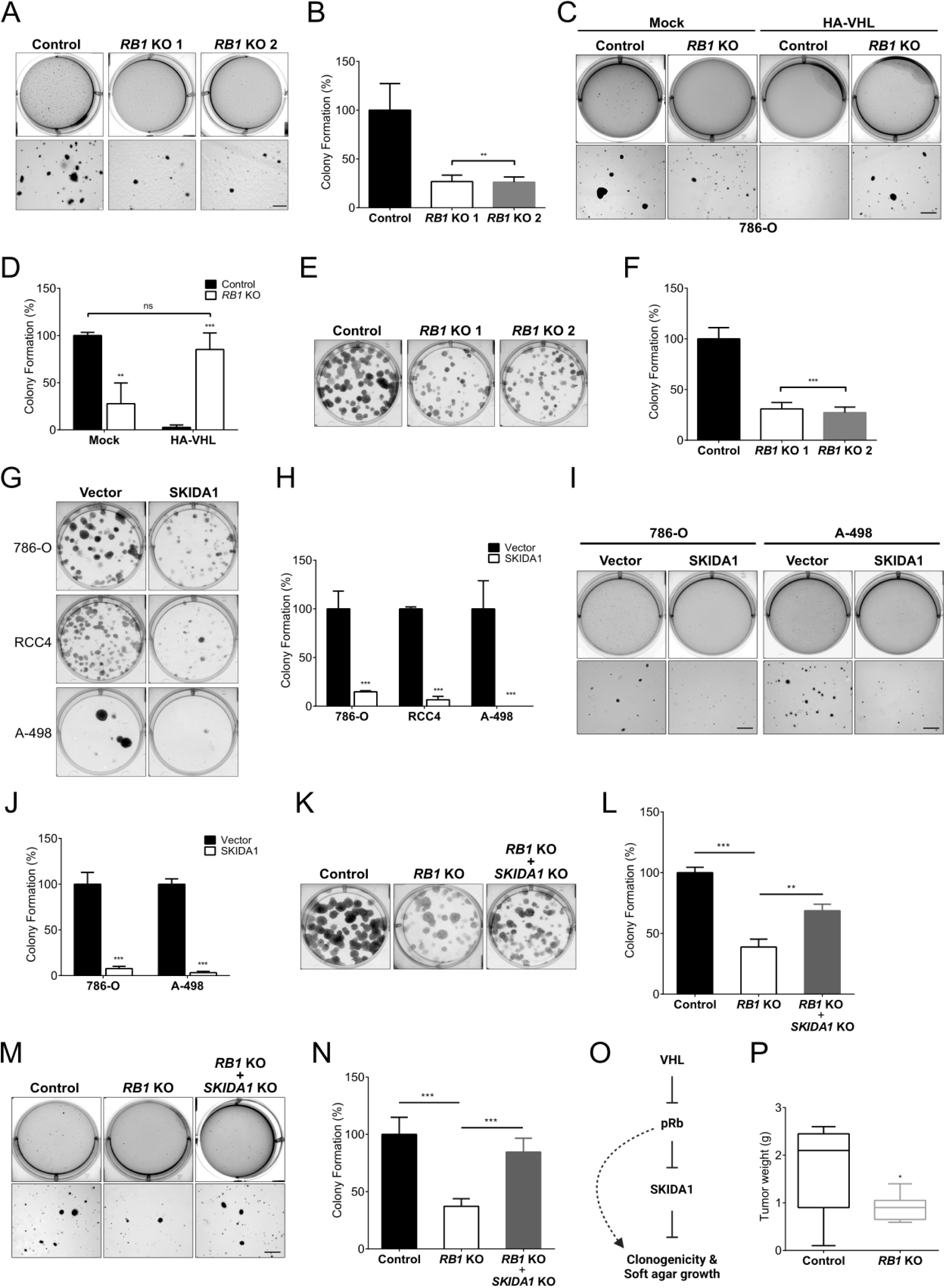
pRb hyperstabilization promotes ccRCC oncogenesis. A. Representative images showing anchorage-independent growth of 786-O control and *RB1* KO cells in soft agar. Scale bar = 500 µm B. Quantification of the number of colonies formed in (A). Colonies were counted under a light microscope. Statistical significance was calculated using one-way ANOVA and Dunnett’s test. C. Representative images showing anchorage-independent growth of 786-O control and *RB1* KO cells stably transfected with either control vector (Mock) or HA-tagged VHL. Scale bar = 500 µm. D. Quantification of the number of colonies formed in (C). Colonies were counted under a light microscope. Statistical significance was calculated using two-way ANOVA and Tukey’s HSD test. E. Representative images showing clonogenic outgrowth of 786-O control and *RB1* KO cells. F. Quantification of the number of colonies in (E). Statistical significance was calculated using one-way ANOVA and Dunnett’s test. G. Representative images showing clonogenic outgrowth of 786-O, RCC4 and A-498 cells stably infected with lentivirus encoding either control plasmid (Vector) or SKIDA1 cDNA containing plasmid. H. Quantification of the number of colonies in (G). Statistical significance was calculated using two-way ANOVA and Fisher’s LSD test. I. Representative images showing anchorage-independent growth of 786-O and A-498 cells stably infected with lentivirus encoding either control plasmid (Vector) or SKIDA1 cDNA containing plasmid. Scale bar = 500 µm. J. Quantification of the number of colonies formed in (I). Colonies were counted under a light microscope. Statistical significance was calculated using two-way ANOVA and Fisher’s LSD test. K. Representative images showing clonogenic outgrowth of 786-O control, *RB1* KO and *RB1*/*SKIDA1* double KO cells. L. Quantification of the number of colonies in (K). Statistical significance was calculated using one-way ANOVA and Tukey’s HSD test. M. Representative images showing soft agar growth of 786-O control, *RB1* KO and *RB1*/*SKIDA1* double KO cells. Scale bar = 500 µm. N. Quantification of the number of colonies formed in (M). Colonies were counted under a light microscope. Statistical significance was calculated using one-way ANOVA and Tukey’s HSD test. O. Schematic illustration showing the VHL-pRb-SKIDA1 signalling axis and effect on clonogenicity and soft agar growth. The dotted line indicates indirect regulation. P. Box and whisker plot showing quantification of weights of tumors formed by 786-O control and *RB1* KO cells subcutaneously injected into the flanks of NOD scid gamma (NSG) mice, via a tumor xenograft assay. 5 injections were performed using control cells and 9 injections using *RB1* KO cells. Statistical significance was calculated using unpaired t-test. (A-P) * p < 0.05, ** p < 0.01, *** p < 0.001. ‘ns’ denotes not significant. Error bars represent standard deviation.

We next assessed the clonogenic capacity of 786-O control and *RB1* KO cells by performing a clonogenic assay, a survival assay based on the ability of a single cell to grow into a colony. Consistent with our findings from the soft agar assay, we observed a significant decrease in clonogenic outgrowth following pRb depletion (Figure 5E-F). Together, these results indicate that pRb is required to maintain the tumorigenic potential of ccRCC cells. We next looked downstream at the potential impact of SKIDA1 on ccRCC clonogenicity and anchorage-independent growth. We found that upregulation of the pRb target SKIDA1 in ccRCC cells led to a significant decrease in clonogenic outgrowth (Figure 5G-H) and anchorage-independent growth in soft agar (Figure 5I-J). Furthermore, depletion of SKIDA1 in 786-O *RB1* KO cells resulted in partial rescue of colony formation (Figure 5K-L) and anchorage-independent growth in soft agar (Figure 5M-N). These results further emphasize that SKIDA1 may be an important downstream tumor suppressor target of the VHL-pRb pathway in ccRCC (Figure 5O). To examine the effect of *RB1* knockout on ccRCC tumor growth *in vivo*, we injected 786-O control and *RB1* KO cells subcutaneously into the flanks of NOD SCID gamma (NSG) mice. All mice were euthanized when the first tumor reached endpoint. Tumors were then excised and weighed. We found that tumors formed by 786-O control cells were significantly larger compared to those formed by *RB1* KO cells, despite one 786-O control injection that failed to grow (Figure 5P). Collectively, our findings indicate that pRb hyperstabilization promotes ccRCC tumorigenesis, likely through transcriptional remodelling including the identified target SKIDA1.

## Discussion

The incidence rate of ccRCC is 8.01 per 100,000 population in patients at least 40 years of age^94^. ccRCC is rarely diagnosed at early stages and tumors at advanced stages are extremely resistant to chemotherapy and radiation therapy^95^. While combination therapy with immune checkpoint inhibitors and anti-angiogenic drugs show improved antitumor efficacy, issues relating to systemic toxicity and drug resistance persist, necessitating research into novel therapeutically tractable pathways^39,96^. Using an unbiased screen to discover novel targets of VHL, we identified pRb as a VHL substrate that is regulated by ubiquitination and subsequent proteasomal degradation. pRb depletion led to decreased clonogenicity and anchorage-independent growth of ccRCC cells. Loss of pRb also led to reduced ccRCC tumorigenesis *in vivo* using mouse xenograft models, highlighting the tumorigenic activity of pRb in ccRCC. Transcriptomic analysis identified SKIDA1 as an important downstream target of the VHL-pRb pathway that is repressed in ccRCC. Overexpression of SKIDA1 significantly decreased ccRCC clonogenicity and anchorage-independent growth. This supports further characterization of the mechanisms of SKIDA1’s tumor suppressive function in ccRCC and an expanded characterization of other potentially important transcriptional targets repressed by pRb upon VHL loss. In this paper, the characterization of pRb hyperstabilization in ccRCC provides a sound rationale for molecular targeting of pRb and its disease-related targets such as SKIDA1.

Although pRb is best known as a tumor suppressor, in this study, we describe a context-specific potential for this protein to act as an oncogene when dysregulated by VHL-deficiency. *RB1* is not a common target for inactivating mutations in ccRCC, so in theory, pRb should be able suppress cell cycle. However, this function is mediated by the unphosphorylated form of pRb and in ccRCC, pRb is inactivated by hyperphosphorylation^55,97,98^ The inactivation of the tumor suppressive function of wild type pRb in ccRCC is likely due to the reported cyclin D1 overexpression, fueled by HIF activation, as well as the inactivation of CDK inhibitors, leading to phosphorylation-based inactivation of cell cycle control^62,99^. Hyperphosphorylated pRb likely promotes tumorigenesis through mechanisms independent of cell cycle. Several studies have described cell-cycle-independent functions of pRb. For example, studies on homozygous *RB1* deletion in mouse embryos showed massive cell death in several tissues including the eye lens, nervous system, and skeletal muscle^72,87,100^. These *RB1* mutant embryos also fail to reach term. Furthermore, pRb could promote resistance to apoptosis induced by ceramide, a lipid second messenger^101^. Our findings also highlight a context-dependent role for pRb in ccRCC whereby pRb functions as an oncogene in the absence of VHL, and as a tumor suppressor in the presence of VHL. As such, targeting of pRb in ccRCC tumors may be restricted to tumors that lack functional VHL expression. Furthermore, the HIF-dependent activity of pRb in ccRCC raises important questions as to the role of pRb in hypoxia regulation in ccRCC. Hence, the efficacy of pRb targeting in ccRCC may be guided by HIF regulation in these tumors.

The function of SKIDA1 is relatively understudied. It was found to be differentially expressed in the regulatory T cells of Autoimmune polyendocrine syndrome type I (APS-1) patients compared to healthy patients^102^. SKIDA1 was also described as the best predictor of *KMT2A* (Mixed-lineage-leukemia) gene rearrangements^103^. Its role however, in tumor development has not been defined. The understudied nature of SKIDA1 and its prognostic value in ccRCC patient survival made it a particularly interesting target to follow up on. Here we describe a role for SKIDA1 in inhibiting ccRCC tumorigenesis by promoting cell death. Mechanistically, SKIDA1 repression in ccRCC may be mediated by the pRb-E2F1 repressive complex as depletion of either pRb or E2F1 rescues SKIDA1 expression in ccRCC cells. This supports previous findings that pRb can regulate E2F1-driven transcription of apoptotic genes^54,104,105^. While our experiments point towards transcriptional regulation of SKIDA1 by the pRb-E2F1 complex, chromatin immunoprecipitation (ChIP) experiments will be needed to validate direct regulation of SKIDA1 by these transcription factors. In this study, we examine VHL-pRb regulation in the context of disease. However, we noted the oxygen-sensitive regulation of pRb by VHL, suggesting that this regulation may play an important role in the cellular adaptation to hypoxia. Further characterization of the VHL-pRb signaling pathway in normal cells may promote our understanding of the cellular hypoxia response and the interplay with disease development. Overall, our findings highlight pRb as a novel target for VHL-mediated degradation and the VHL-pRb-SKIDA1 pathway as a potential therapeutic avenue for ccRCC treatment.

## Acknowledgements

The authors acknowledge the support from CIHR grant #153034 (R.C.R), Cancer Research Society Grants # 840178 & # 863801(R.C.R), and Natural Sciences and Engineering Research Council of Canada #2023-05587(R.C.R). M.K. is a recipient of the Asan Foundation PhD fellowship. The authors also acknowledge the following Core facilities from the University of Ottawa and the Ottawa Hospital Research Institute (OHRI) for use of their facility, equipment, and expertise: the Cell Biology and Imaging Acquisition Core (RRID:SCR_021845), the Flow Cytometry and Virometry Core (RRID:SCR_023306), the Genome Engineering and Molecular Biology Core (RRID:SCR_022954) and the OHRI StemCore Laboratories (RRID:SCR_012601).

## Materials and methods

### Cell Culture and Reagents

All cell lines used in cell culture were purchased from the American Type Culture Collection (ATCC). 786-O, 769-P and RCC4 cells were grown in RPMI 1640 medium (350-000-CL, Wisent) supplemented with 10% (vol/vol) fetal bovine serum (FBS). A-498, 293A and U2OS cells were grown in Dulbelcco’s modified Eagle’s medium (319-015-CL, Wisent) supplemented with 10% (vol/vol) FBS. All cells were maintained in a humidified incubator at 37°C, 5% CO_2_, and 21% O_2_ (normoxia). Assays were performed on arrested cultures grown to confluence in order to control for pRb cell cycle function. For hypoxic treatments, cells were placed in a hypoxia chamber at 37°C, 5% CO_2_, and 0.5% O_2_ for 24 hours. MG132 (Peptides International) was used at 10µM.

### Western Blot and Antibodies

Whole cell lysates were prepared by lysing cells in 1x Laemmli buffer. After boiling at 95°C for 10 minutes, samples were separated by SDS-PAGE and transferred on to polyvinylidene fluoride (PVDF) membranes. After blocking, membranes were incubated in primary antibody diluted in 5% bovine serum albumin (BSA) in TBST. Membranes were washed with TBST and incubated with the appropriate horseradish peroxidase (HRP)-conjugated secondary antibody diluted in 2% non-fat milk in TBST. After washing, blots were developed with the chemiluminescence method and imaged using the Bio-Rad ChemiDoc imaging system.

Antibodies against pRb (9313), HA (2999), VHL (68547), and Ubiquitin (3936) were from Cell Signaling Technology. Antibodies against Vinculin (V9131), β-actin (A5441), α-tubulin (T6199) and Flag (A8592) were from Sigma Aldrich. Anti-HIF2α (NB100-122) antibody was from Novus Biologicals. Glut-1 (115730) antibody was from Abcam. Anti-SKIDA1 (ARP69749_P050) antibody was from Aviva systems biology. Peroxidase-conjugated sheep anti-mouse (NB1206808) and donkey anti-rabbit (NB7185) antibodies were purchased from Novus Biologicals. Antibodies used for co-immunoprecipitation include mouse anti-HA (homemade), anti-Flag M2 (A2220, Sigma) and mouse anti-pRb (G3-245, BD Biosciences).

### Quantitative real-time polymerase chain reaction (qRT-PCR)

Total RNA was extracted from cells using the PureLink RNA Mini Kit (12183018A, Invitrogen) according to the provided protocol. On-column genomic DNA digestion was performed using the PureLink DNase Set (12185010, Invitrogen). First strand cDNA synthesis was performed with the iScript cDNA Synthesis Kit (#1708891, Biorad) according to provided protocol. Real-time quantititative PCR (qRT-PCR) detection was performed using the Luna® Universal qPCR Master Mix (M3003, New England Biolabs) and the CFX96 Real-Time PCR System. 10 ng of cDNA was used as input for qRT-PCR reactions. Real-time PCR was performed in triplicate and relative gene expression was calculated using the 2^-ΔΔCt^ method^106^ following normalization of Ct values to a GADPH control. Gene-specific primers were designed using the IDT PrimerQuest tool.

Real-time PCR primers are listed below:

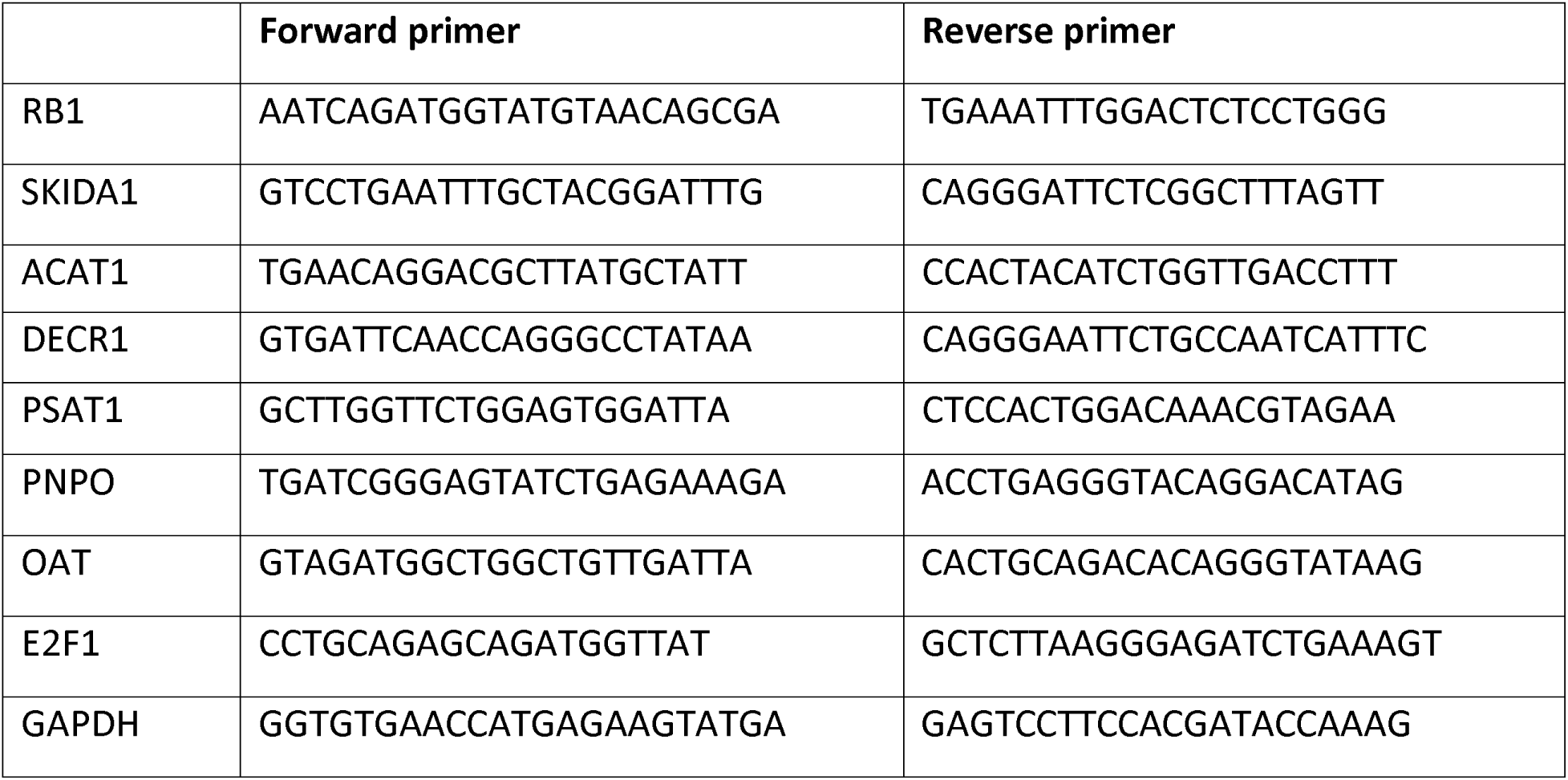

### Virus Production and transduction

HEK293T cells were used for lentivirus production. The packaging plasmids psPAX2 (#12260) and pMD2.G (#12259) were purchased from Addgene. Viruses were harvested at 48 h and 72 h post-transfection, filtered through 0.45μm PVDF filter and immediately used to infect target cells. The virus-containing media was supplemented with 8 µg/mL polybrene to improve transduction efficiency. Transduced cells were then selected for using the appropriate antibiotic. Stable cell lines were created by selecting for viral integration using 1 µg/ml puromycin or 0.5 mg/ml G418, as appropriate.

### Transfection and Co-immunoprecipitation assays

Plasmid transfection of 293A/T cells was performed using polyethylenimine (PEI) (23966-1, Polyscience) reagent 48 hours prior to lysis. Plasmid transfection of 786-O cells was achieved by electroporation using the Amaxa Nucleofector 2b device and manufacturer recommended conditions. siRNAs were transfected with Lipofectamine RNAiMAX transfection reagent (13778030, Thermo Fisher Scientific).

For immunoprecipitation, cells were harvested in mild lysis buffer (10mM Tris pH 7.5, 100mM NaCl, 10mM EDTA, 50mM NaF, 1% NP-40) supplemented with complete protease inhibitor. Lysates were clarified by centrifugation, and then incubated with the appropriate antibody and beads for 1 hour. Bound complexes were washed with mild lysis buffer and eluted by boiling in 1x Laemmli buffer at 95°C for 10 minutes. Bound proteins were then resolved using SDS-PAGE followed by western blot analysis. For denaturing immunoprecipitation, after clarification of cell lysate, sodium dodecyl sulphate (SDS) was added to a final concentration of 1% and the lysate was boiled at 95°C for 5 minutes in order to denature proteins and disrupt protein-protein interactions. The cooled lysate was then diluted to a final concentration of 0.1% SDS prior to immunoprecipitation.

### BioID assay

Cells transfected with VHL-Flag-BirA plasmid were incubated with 50 μM biotin for 1 hour. Cells were lysed with high salt RIPA buffer and snap-frozen in liquid nitrogen. Lysates were thawed rapidly in a 37°C water bath and clarified by sonication followed by centrifugation. Lysates were pre-cleared using protein G agarose beads (16-266, Sigma Aldrich). The pre-cleared lysates were incubated with streptavidin-agarose beads (20359, Thermo Fisher Scientific) to purify biotinylated proteins. Biotinylated proteins were then eluted off the beads. The samples were resolved on a NuPAGE 10% Bis-Tris gel (NP0315BOX, Thermo Fisher Scientific) and visualized using the SimplyBlue SafeStain (465034, Thermo Fisher Scientific). Bands were then excised, trypsin digested, and analyzed using mass spectrometry.

### Live/Dead cell staining

Live/dead cell staining was performed using the ReadyProbes® Cell Viability Imaging Kit (Blue/Green) (R37609, Thermo Fisher Scientific). The appropriate volume of each dye was added directly to cells and incubated for 15 minutes at room temperature. The NucBlue® Live reagent stains all nuclei whereas the NucGreen® Dead reagent stains only the nuclei of cells with compromised plasma membrane integrity. Images were obtained using the EVOS M5000 microscope at 10x objective. The Fiji software was used for image analysis and adjustment of image parameters.

### TUNEL assay

Adherent cells were rinsed twice with PBS and fixed with 4% paraformaldehyde (PFA) for 1 hour at 25°C. Cells were rinsed with PBS, then permeabilized with 0.1% Triton X-100 in 0.1% sodium citrate for 2 minutes on ice. Cells were rinsed twice with PBS. 50 µl TUNEL reaction mixture (11684795910, Sigma Aldrich) was added to cell monolayer and incubated for 60 minutes at 37°C in a humidified atmosphere in the dark.LLCells were then rinsed 3 times with PBS and counterstained with 1 µg/ml 4L,6-diamidino-2-phenylindole (DAPI). For negative control, cells were incubated in 50 µl/well Label Solution (without terminal transferase) instead of TUNEL reaction mixture. For positive control, cells were incubated with DNase I (3 U/ml in 50 mM Tris-HCl pH 7.5, 10 mM MgCl_2_, 1 mg/ml BSA) for 10 minutes at 25°C to induce DNA strand breaks prior to labeling procedures. Images were obtained using the EVOS M5000 microscope at 10x objective. The Fiji software was used for image analysis and adjustment of image parameters.

### Sulforhodamine B (SRB) assay

Cells were fixed in media containing 10% trichloroacetic acid (TCA) (TB0968, Bio Basic) at 4°C for 2 hours. Following wash off of fixation solution, cells were stained with 0.04% (wt/vol) Sulforhodamine B sodium salt (HY-D0974, MedChemExpress) for 30 minutes. Cells were rinsed with 1% (vol/vol) acetic acid to remove unbound dye. 150 µl of 10 mM Tris base solution (pH 10.5) was then added to solubilize the protein-bound dye. Dye absorbance (570 nm) and background absorbance (650 nm) were obtained using a microplate reader. Final absorbance reading was obtained by subtracting background absorbance from dye absorbance.

### Soft-agar colony formation assay

The soft agar colony formation assay was done on 6-well plates. The bottom layers consisted of 1.5 mL of 1% low melting point agarose (16520050, Thermo Fisher Scientific) in 1X RPMI media supplemented with FBS. The upper layers consisted of 7500 cells embedded in 1.5 mL of 0.5% agarose in 1X RPMI media supplemented with FBS. The layers were allowed to solidify for 5 minutes at 4°C, then placed in a 37°C incubator for the remainder of the assay. Every 3 days, 200 µL of complete media were added onto the semi-solid media to prevent desiccation. After 5 weeks, colonies were stained with 100 µg/mL iodonitrotetrazolium chloride (IB0280, Bio Basic) solution. Colonies were counted under a light microscope. Colonies greater than 100 µm in diameter were quantified. Images were obtained using the Bio-Rad ChemiDoc imaging system and the EVOS M5000 microscope.

### Clonogenic assay

150 cells were seeded on a 6-well plate and cultured for 10 days. Cell culture media was replaced every 2-3 days. For drug treatments coupled with clonogenic assay, 150 cells were seeded on a 6-well plate a day before drug treatment. Drugs treatments were done for 3 days, washed off once with phosphate-buffered saline (PBS), and replaced with regular media for an additional 7 days. Following colony formation, cells were rinsed with PBS, fixed with 3 vol methanol + 1 vol acetic acid for 5 minutes and stained with 0.5% (wt/vol) crystal violet in methanol for 15 minutes. The dye was rinsed off with tap water until background was clear. Pictures of the visible colonies were obtained using the Bio-Rad ChemiDoc imaging system.

### RNA sequencing and data analysis

Total RNA from three biological replicates was extracted using Trizol reagent (15596026, Thermo Fisher Scientific). RNA was initially quality controlled by Qubit and Tapestation. Libraries were prepared using the Truseq kit from Illumina using the manufacturer’s protocol (https://www.illumina.com/products/by-type/sequencing-kits/library-prep-kits/truseq-stranded-mrna.html). Quantity and quality of libraries were assessed by qPCR and LabChip, respectively. All libraries were sequenced on 1 lane of NovaSeq S1 (PE100), to generate 50 millions read per condition.

Raw sequencing data was processed using GenPipes^107^ with standard settings, which aligns reads to GRCh37 (hg19) with *STAR* 2-passes mode and counted to gene features using ‘htseq-count’ function from *HTSeq*. The raw count data was normalized and analyzed by *DESeq2*. The differentially expressed genes (DEG) (adjusted p-value < 0.05, fold change > 1.5 or > 2.0) were compared to HIF2α-activated genes (downloaded from GSE149005 and re-analyzed by *limma* for significant DEGs with adjusted p-value < 0.05), downregulated genes in ccRCC versus normal kidney tissues (downloaded from the International Cancer Genome Consortium (ICGC) and analyzed by DESeq2), and E2F1-regulated genes (downloaded from TRANSFAC database from ChIP-X enrichment analysis^82,108^). The comparisons between the DEGs in each group were visualized using *ggvenn*, and the pathway enrichment analysis was done using *pathfindR*. The enriched pathways were filtered based on biological relevance and visualized in a dot plot.

### Tumor Xenograft Assay

For tumor xenografts, NOD scid gamma mice at 6-8 weeks of age and 20 – 23 g average body weight were used. The mice were housed and maintained in laminar flow rooms under specific pathogen-free conditions. All animal procedures were performed according to the guidelines of the Canadian Council on Animal Care (CACC). The protocol for animal studies was approved by the Animal Care Committee of the University of Ottawa.

Briefly, 786-O control and *RB1*-KO cells were harvested at exponential growth phase using 0.25% trypsin-EDTA (25200-056, Thermo Fisher Scientific). Cells were washed once and resuspended in PBS. The number of viable cells was determined by trypan blue exclusion assay. 8 million viable cells were resuspended in 50% (vol/vol) Matrigel to a final volume of 100 µL and injected subcutaneously into the flank of each mouse. The injection sites were manually palpated twice weekly until tumors were established. Caliper measurements of the length and width of the tumor were taken weekly until the onset of tumor growth and then twice weekly until endpoint. Mouse weights were recorded every 3 days. All mice were euthanized at 11 weeks post-injection when the first mice had reached endpoint. At the end of the experiment, tumors from each group were excised and weighed.

### CRISPR/Cas9 Genome Editing

Monoclonal 786-O *RB1*-KO cell lines were generated with CRISPR/Cas9 technology. 786-O cells were transfected with PX458 plasmid (#48138, Addgene) containing *RB1* targeting single guide RNAs (sgRNAs). Cells containing the plasmid of interest were selected by single cell sorting of green fluorescent protein (GFP)-expressing cells. Monoclonal cells were screened for the gene knockout by western blot and later, Sanger sequencing of the region of interest. Successful knockout clones were then selected and expanded.

The following guide RNAs were used to create monoclonal *RB1* knockout cell lines:

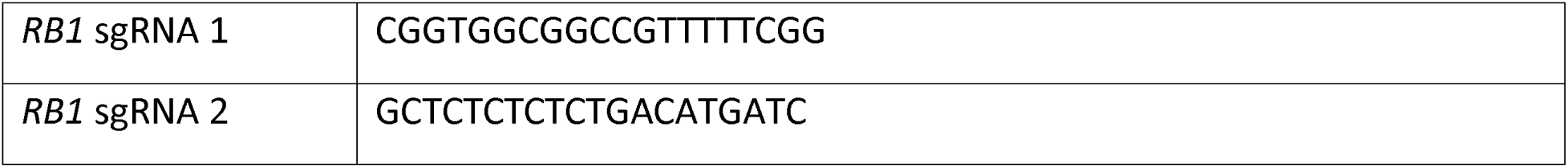

Polyclonal knockout cell lines were generated by stable expression of Cas9 (#52962, Addgene) in target cells, followed by transduction with lentivirus containing the corresponding guide RNAs in the pCLIP-dual-SFFV-ZsGreen vector backbone. Each pCLIP-dual-SFFV-ZsGreen vector contains 2 guides (gRNA_a and gRNA_b) targeting the gene of interest. All guides were designed against human genes. pCLIP-dual-SFFV-ZsGreen sgRNAs were obtained from transEDIT-dual CRISPR Whole Genome Arrayed Library from Transomic technologies. Target sequences were as follows:

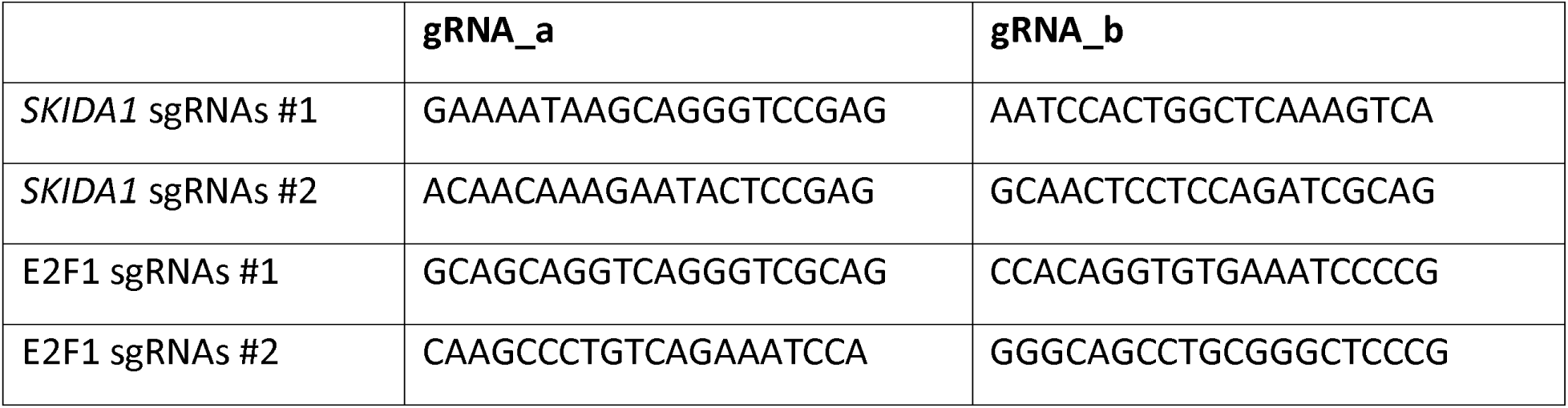

### Plasmids and siRNA constructs

HA-VHL wt-pBabe-puro (#19234, Addgene) plasmid was used for transient transfections. HA-VHL-pRc/CMV (#19999, Addgene) plasmid was used to create stable cell lines. Flag-pRb plasmid was obtained by cloning Flag-tagged pRb into the pLVX-M-puro vector (#125839) from Addgene. For overexpression studies, the gene of interest was cloned into the pLVX-M-puro vector using Gibson cloning. All plasmids were sequenced to confirm validity. ON-TARGETplus Non-targeting Control siRNAs (D-001810-01-05) and siGENOME Human RB1 siRNA smartpool (M-003296-03-0005) were purchased from Dharmacon.

### Statistical analysis

Statistical analysis was performed using the GraphPad Prism6 software. Data are presented as mean ± standard deviation (SD) unless otherwise indicated. p-values < 0.05 were considered statistically significant. * p < 0.05, ** p < 0.01, *** p < 0.001. ‘ns’ denotes not significant.

