## Supplementary figures and images for "Loss of VHL-mediated pRb regulation promotes clear cell renal cell carcinoma"

### Supplemental Figure 1

S1

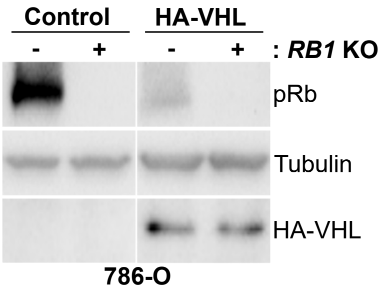

### Supplemental Figure 2

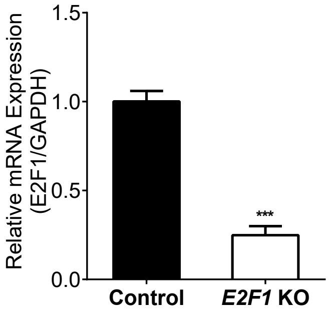

### Supplemental Figure 3

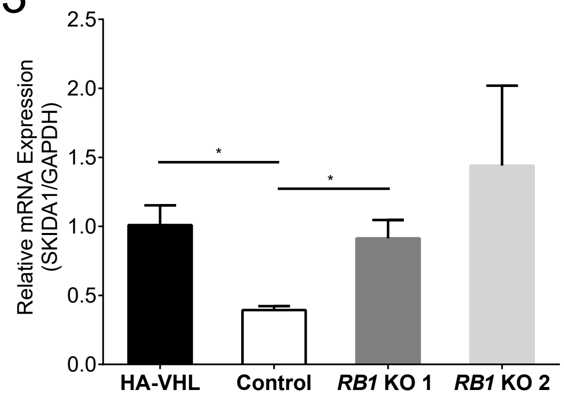

### Supplemental Figure 4

S4

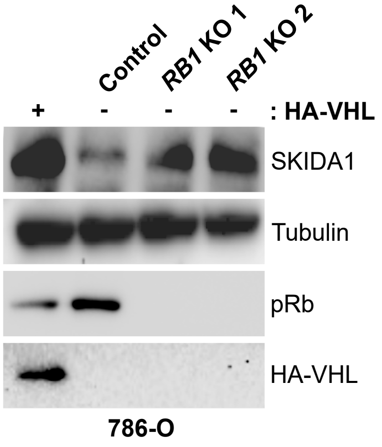

### Supplemental Figure 5

# S5

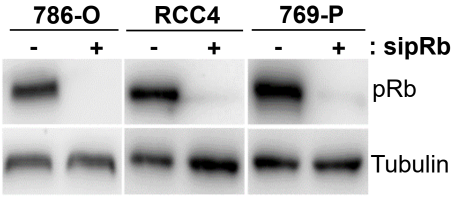

### Supplemental Figure 6

S6

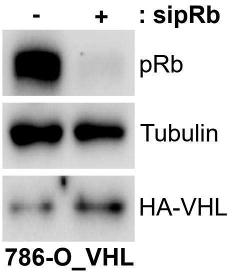

### Supplemental Figure 7

# S7

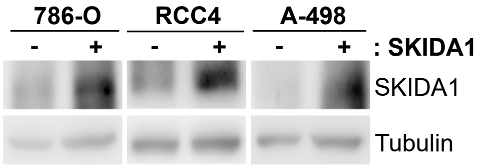

### Supplemental Figure 8

S8

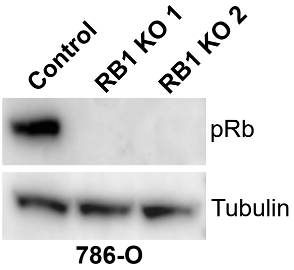
