## Supplemental Figure 9 for "Loss of VHL-mediated pRb regulation promotes clear cell renal cell carcinoma"

# S9

### 786-O *RB1* KO 1

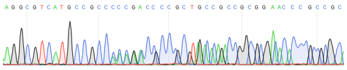

| | |
|---|---|
| TCATGCCGCCCAAAACCCCCCGAAAAACGGCCGCCACCGCCGCCGC | WT |
| TCATGCCGCCCAAAACCCCCCGAAAAA-GGCCGCCACCGCCGCCGC | Allele 1 |
| TCATGCCGCC-------------------------ACCGCCGCCGC | Allele 2 |

### 786-O *RB1* KO 2

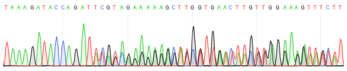

| | |
|---|---|
| AAAGATACCAGATCATGTCAGAGAGAGAGC | WT |
| AAAGATACCAGAT-ATGTCAGAGAGAGAGC | Allele 1 |
| AAAGATACCAGAT----TCAGAGAGAGAGC | Allele 2 |
